# Neural correlates of implicit memory for reoccurring sound patterns

**DOI:** 10.64898/2025.12.18.695138

**Authors:** Ylka Kolken, Chengran Li, Vincent van de Ven, Peter De Weerd, Ryszard Auksztulewicz, Roberta Bianco, Maria Chait, Federico De Martino

**Author notes:** Corresponding author: Ylka Kolken, Maastricht University, Maastricht, The Netherlands.

## Abstract

The human brain exhibits remarkable sensitivity to auditory patterns. Previous research has shown that beyond the detection of such patterns, humans form memories for them that are implicit, robust to interference, and persistent over time. Here, we examine the mechanisms underlying implicit memory formation for reoccurring regular tone patterns using UHF fMRI. Prior to scanning, implicit memory was induced through a standard behavioral task where participants detected regular patterns embedded in random tone sequences. Unbeknownst to participants, five regular patterns reoccurred sporadically (every 1.5 min) across trials. Consistent with previous reports, behavioral performance indicated a reaction time advantage to reoccurring patterns, confirming the formation of implicit memory traces. During passive listening in the scanner, regularity engaged the superior temporal gyrus, inferior frontal gyrus, insula, putamen, and hippocampus, a result we interpret as reflecting the higher perceptual precision of regular patterns. Reoccurrence interacted with regularity in the superior temporal gyrus and a frontal cluster, showing higher activity for reoccurring compared to novel regular patterns. The hippocampus showed sensitivity to reoccurrence in both regular and random patterns. Together, these results indicate that the neocortex may selectively amplify reoccurring inputs in line with their enhanced (memory driven) perceptual precision.

## Introduction

Our auditory system is constantly exposed to complex streams of sound. One of its most remarkable, yet ubiquitous abilities is to uncover patterns, often without the listeners’ conscious awareness [1–6].

Previous work has demonstrated that the emergence of a regular pattern within a stream of random tone pips is associated with a gradual increase in sustained (a-periodic, low-frequency) M/EEG activity in the inferior frontal gyrus (IFG), Heschl’s gyrus, and the superior temporal gyrus (STG) [3, 6, 7]. This increase is accompanied by decreased tone-locked responses [6], consistent with the idea that regularity extraction involves both local suppression of predictable inputs, in line with top-down signaling, resulting in attenuated cortical responses to expected stimuli [8, 9], and an increase in the precision of priors for regular stimuli, reflected in heightened activity associated with the growing reliability of predictable sequences.

Interestingly, this putative precision-linked network also involves the hippocampus [3, 6, 7]. The hippocampus has a critical role within the predictive hierarchy [10]. Its subfields perform specialized functions, such as pattern separation and completion, that are compatible with predictive processes [11, 12], and underlie its involvement in detecting sequence statistics [13–15]. Importantly, the observed hippocampal involvement during passive listening underscores its often overlooked contribution to the automatic processing of dynamically unfolding auditory information [16].

When regular patterns reoccur across trials, even sparsely, they become more readily detectable, demonstrating a considerable implicit detection time advantage over novel regular patterns [17, 18]. Modelling using prediction by partial matching (PPM; [19]) demonstrates that this effect can be explained by the progressive strengthening of memory traces associated with experienced tonal transitions (*n*-grams). This strengthening leads to a faster reduction of surprise (or information content) associated with tones in the pattern, thereby facilitating more rapid detection of previously encountered patterns. Alternative explanations, particularly in the context of *memory for noise* findings, suggest sensitivity to specific spectro-temporal idiosyncrasies [1, 20, 21]. These are thought to involve potentiated synapses that enable faster neural responses and more salient detection when the same features reoccur.

Accumulating work demonstrates that this memory effect is present across the age span, robust to interference, and long-lasting [17, 22], indicative of a remarkable human ability to form implicit memory for acoustic patterns [1, 14, 22, 23]. A critical question concerns where such information is stored. Preliminary M/EEG evidence suggests that the detection of reoccurring regularities recruits the same auditory–frontal network involved in the extraction of novel regular patterns [7].

Here, we use ultra high field (UHF) 7-Tesla functional magnetic resonance imaging (fMRI) to investigate how the brain encodes and retains brief structured regular sound sequences embedded in a stream of random sound sequences. To do so, participants were trained before the scanning session to detect regularities within random sound sequences, in which, unbeknownst to the participants, a subset of regular sequences reoccurred. In the immediately following fMRI session, participants passively listened to ongoing sound sequences containing novel and reoccurring regularities.

## Results

### Behavioural Results

Fourteen participants listened to 5-second sequences of contiguous 50ms tone pips and pressed a key when they detected the emergence of a regularity from a random sequence of tones (50% of trials). Presented sequences were either random throughout (RAN; tones uniformly sampled from a pool of 20 values) or transitioned into a regular structure (REG), which consisted of a repeating 20 tone (1 s long cycle) pattern (Figure 1A). REG sequences were either novel (REGn) or reoccurring (REGr). The stimulus set also included control stimuli (STEP and CONT) with which we estimated listeners’ basic reaction time to a simple sound change. To estimate the regularity detection latency, we subtracted response times (RTs) to STEP from RTs to the REG conditions. Each participant completed five blocks, each lasting approximately eight minutes.

**Figure 1:**
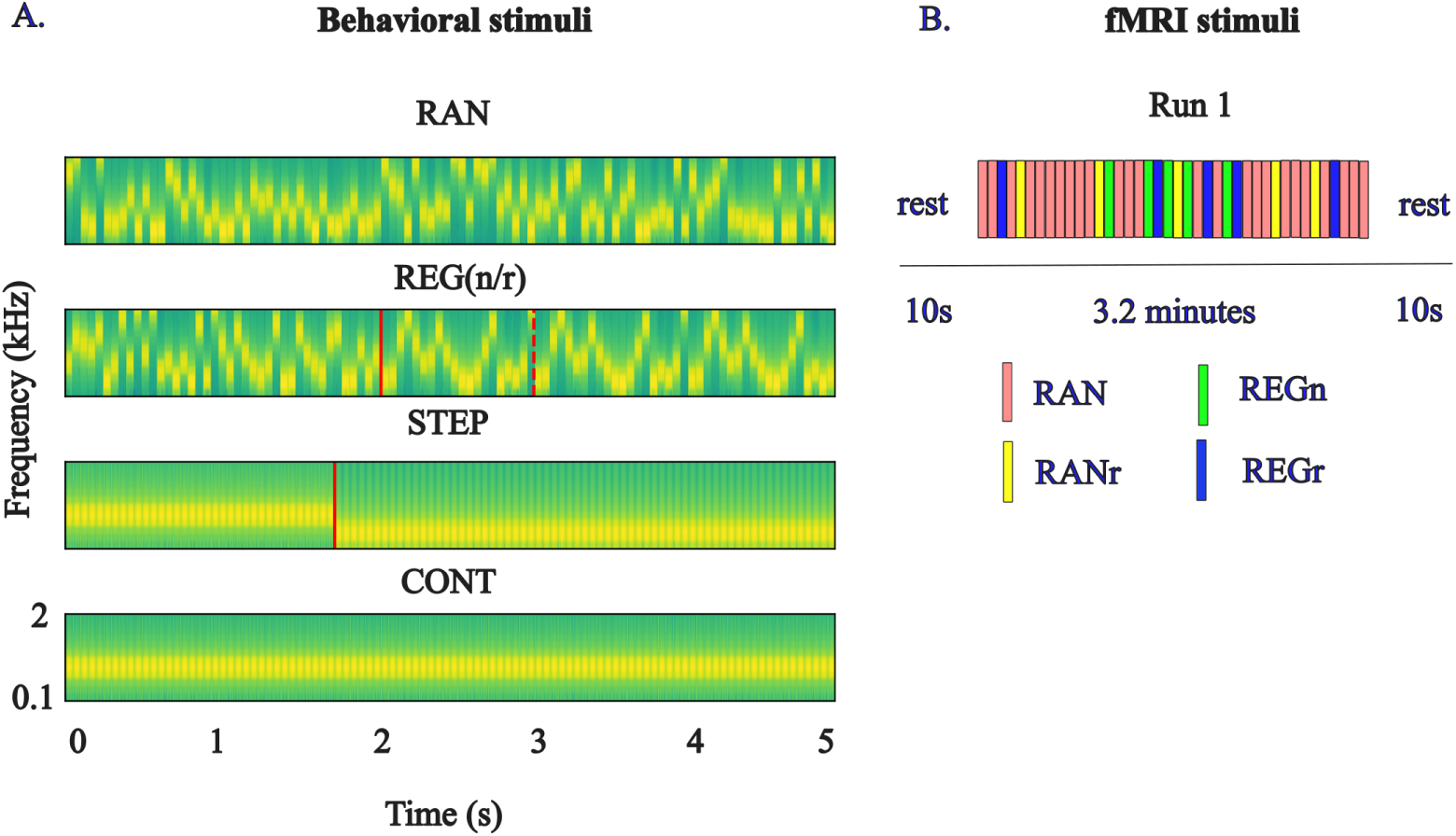
Stimuli used in the behavioural (A) and fMRI session (B). A. Spectrograms of example stimuli: RAN sequences were random sequences of tones; REG sequences contained a transition (indicated with a red line) form RAN, to a regularly repeating pattern. REG patterns were either novel (REGn) or reoccurring (REGr). The red dashed line marks the point at which the pattern starts repeating and hence becomes statistically detectable. During the behavioural session, data were acquired in five blocks. Each block lasted approximately eight minutes containing 30 REGr sequences (five unique sequences reoccurring six times), 30 REGn, 30 RAN, 5 STEP, and 5 CONT sequences. B. After the behavioural session, participants took part in an fMRI session (passive listening). The same stimuli used in the behavioural session were presented as a continuous stream of random tones (RAN) with occasional embedded bursts of REGn, REGr (the same as in the behavioural session) and RANr patterns. The latter were added to control for the mere repetition effect (see text). The figure shows an example of one half of an fMRI run. Stimuli were presented as a continuous RAN sequence, into which occasional 2-second bursts of REGn (5), REGr (5 – identical to those in the behavioural session), and RANr (5) were embedded. In the figure, every burst of REGr, REGn, or RANr is always preceded by 3000 ms of RAN.

Participants’ sensitivity to regularity (d’) plateaued at near ceiling performance after the first block (Figure 2A). The difference in RTs between reoccurring regularities (REGr) and novel regularities (REGn) changed significantly across training blocks (block by condition interaction - F(4,65)=4.218, p=.004, *η*^2^ = 0.206). Further analysis revealed that, already in block 2, the RTs were significantly different between REGr and REGn (t(13)=5.957, p=<.001, d=1.592) (Figure 2B and D). This indicates that participants implicitly learned the reoccurring regularities.

**Figure 2:**
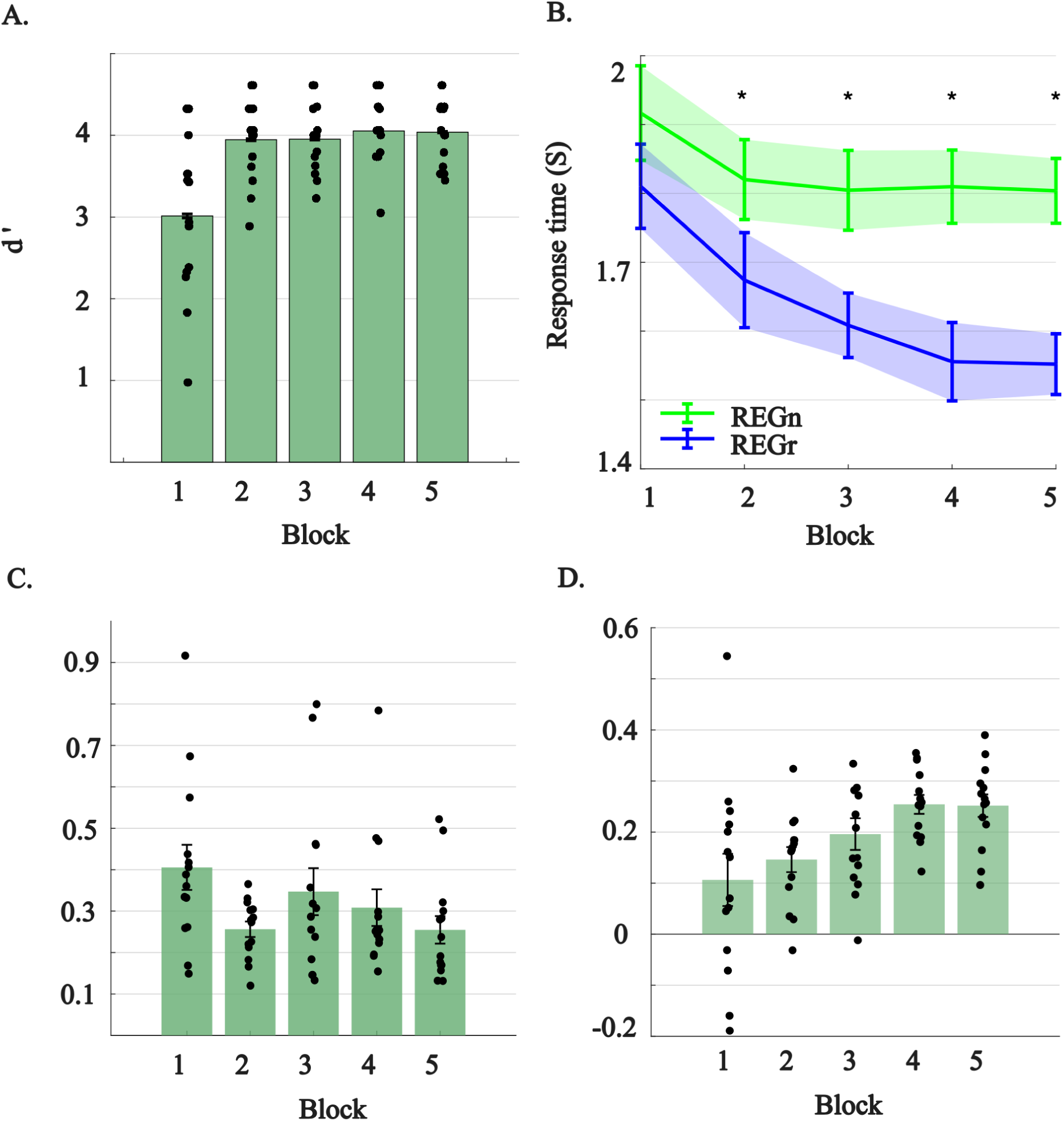
A. Participants’ sensitivity to regularity (d’) across experimental blocks B. Response times across blocks. Note that these RTs are baseline corrected (the RTs to the STEP condition in each block were subtracted from both the RTs of REGr and REGn). Error bars represent SEM C. RTs to STEP. D. Average and individual response time advantage (RTA - [RT REGn - RT REGr]) reflecting memory formation for reoccurring regularities.

### fMRI results

In the 7T fMRI session, the same participants passively listened to continuous streams of sound sequences. Stimuli were presented as a continuous RAN sequence, into which occasional 3-second bursts of REGn, REGr (the same as presented in the behavioural session), and RANr were embedded (Figure 1B). RANr sequences are 5 different 3 second long RAN sequences (chosen randomly for each participant) that reoccurred exactly at the same rate as REGr sequences. They were included to dissociate true memory effects from stimulus repetition effects (see Methods). We collected fMRI data at UHF (7T; TR=1s, 1.6mm isotropic) and, after standard pre-processing, statistical analyses used a fixed-effects GLM with predictors for each condition (REG, REGr, RAN, RANr). Neocortical results were projected to a standard (Talairach) space, hippocampal responses were analysed in native space.

At the group level, sound presentation (t-omnibus test vs. silence) was associated with significantly larger responses in bilateral auditory (Heschl’s gyrus, planum temporale, planum polare, anterior and posterior STG) and frontal cortices (Figure 3 - t(37257)>2.47, q(FDR)<0.05 - individual results of the same analysis are reported in Supplementary figure S1). These results show the reliability of the design in activating relevant areas in the lateral temporal and frontal cortices.

**Figure 3:**
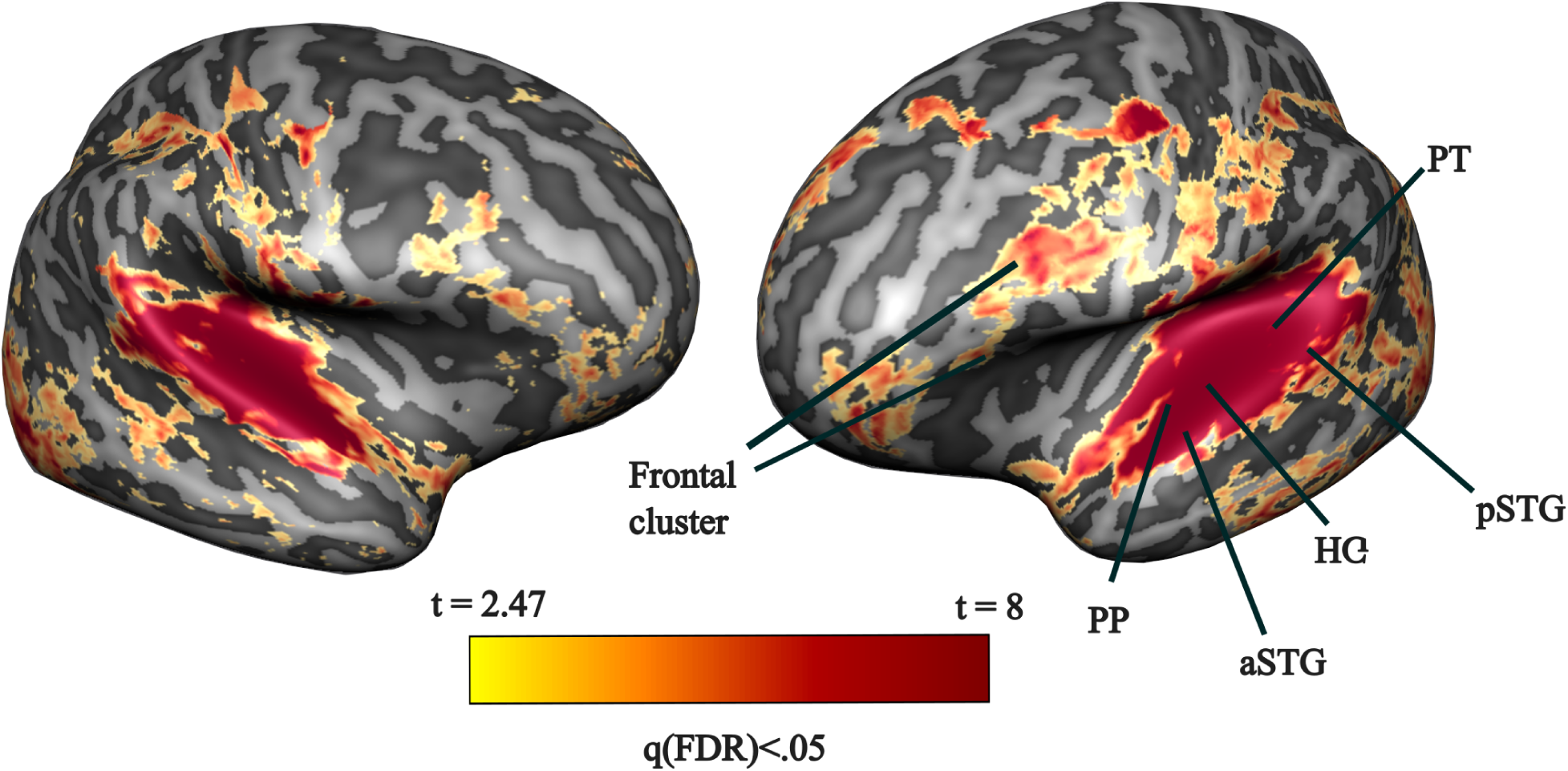
Statistical analysis of the response to any sound (REGn, REGr, RAN, RANr) in our design compared to no sound presentation (t-omnibus test - group level, fixed effects). HG = Heschl’s gyrus, PT = planum temporale, PP = planum polare, pSTG = posterior superior temporal gyrus, aSTG = anterior superior temporal gyrus. The analysis was performed in volumetric space and is here sampled on the average cortical surface of the 14 volunteers. The statistical map was corrected for multiple comparisons using FDR and thresholded at a level of q(FDR)<0.05 (equivalent to a t(37257)>2.47).

For the effect of regularity at the whole brain level, and after correction for multiple comparisons, the presentation of regular vs. random sequences was associated with larger responses in bilateral frontal cortical areas (broadly encompassing the middle/inferior frontal gyrus and anterior insular regions), bilateral (medial and posterior) STG and bilateral putamen (Figure 4 – t(37257)>2.85, q(FDR)<0.05). The IFG and STG have been previously implicated in the detection of statistical regularities in sound sequences, and our findings replicate this pattern [3, 6, 7].

**Figure 4:**
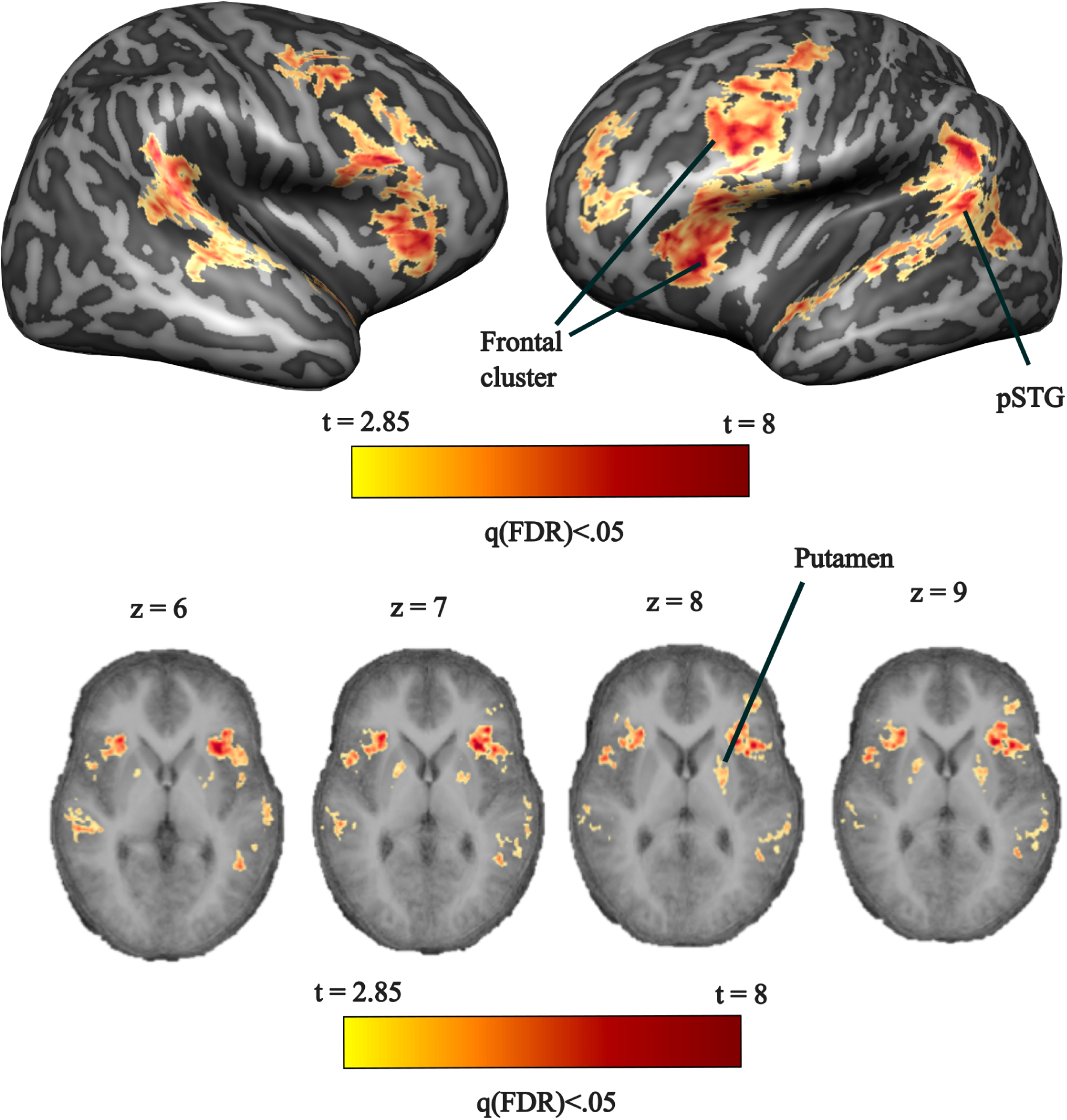
Statistical analysis (t-test) of the effect of regularity (contrasting regular sounds [REGn and REGr] to random patterns [RAN and RANr]) at the group level (fixed effects). pSTG = posterior superior temporal gyrus. The analysis was performed in volumetric space and is also sampled here on the average cortical surface of the 14 volunteers (top) for visualization purposes. The bottom panels present four transversal cuts (from inferior to superior) highlighting the effect in the putamen (bilaterally). The statistical map was corrected for multiple comparisons using FDR and thresholded at a level of q(FDR)<0.05 (equivalent to a t(37257)>2.85).

For the effect of memory, we considered the interaction of reoccurrence and regularity by comparing the difference between reoccurring regular patterns and novel regular patterns (REGr vs. REGn) to the difference between reoccurring random patterns and novel random patterns (RANr vs. RAN). By doing so, we could identify regions where the reoccurrence effect reflected memory for previously learned patterns rather than a simple increase in signal-to-noise ratio (SNR) due to stimulus repetition. A bilateral network comprising the STG and a frontal cluster exhibited a significant interaction (reoccurrence by regularity) with a larger effect of reoccurrence in regular patterns (REGr vs. REGn > RANr vs. RAN; Figure 5; t(37262)>2.97, q(FDR)<0.05). These results indicate that reoccurring regularities engaged a similar network associated with processing regular patterns, and that these regions selectively amplify the response to reoccurring regularities—consistent with a true memory effect. The analysis of the main effect of reoccurrence (contrasting reoccurring regular and random patterns to novel regular and random patterns) did not result in any significant response at the neocortical level.

**Figure 5:**
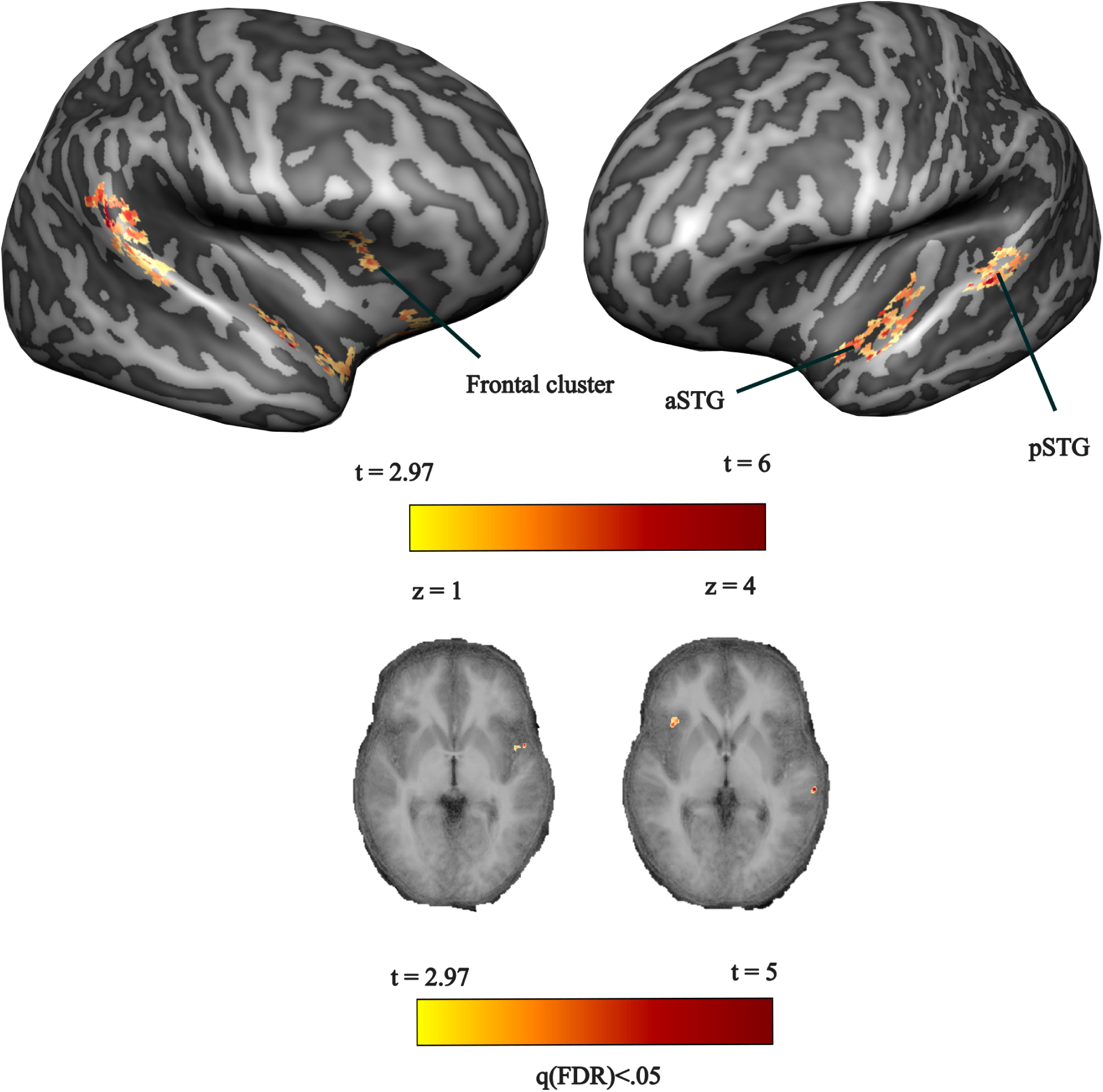
Statistical analysis (t-test) of the interaction between regularity (regular [REG] or random [Ran]) by reoccurrence (reoccurring or novel) at the group level (fixed effects). Positive values indicate a larger reoccurrence effect in the regular patterns than in the random patterns. The analysis is presented using the same conventions as in Figure 4, with effects displayed on the average cortical surface (top). Transversal slices (bottom) show the activation of the frontal cluster and STG.

Moving beyond whole-brain analyses of the neocortical surface, we focused on responses in the hippocampus. Using an automatic pipeline [24], we defined five subfields in the individual bilateral hippocampus (dentate gyrus [DG], subiculum [SUB], cornu ammonis 1-3 [CA1-3] - Supplementary figure S2 displays the segmented regions of interest [ROIs] in the individual anatomy of all participants). We did not separate left and right hippocampi, as we had no specific hypotheses regarding potential hemispheric differences. The effects of regularity were significant in dentate gyrus (t=-3.374; q=0.0019), CA1 (t=-2.658; q=0.0131), and Subiculum (t=-3.993; q=0.0003) (Figure 6A - for all effect sizes and q-values [including non-significant effects] see supplementary Table 1). The estimate of the response was smaller for regular compared to random sounds (light and dark grey in Figure 6A), respectively).

**Figure 6:**
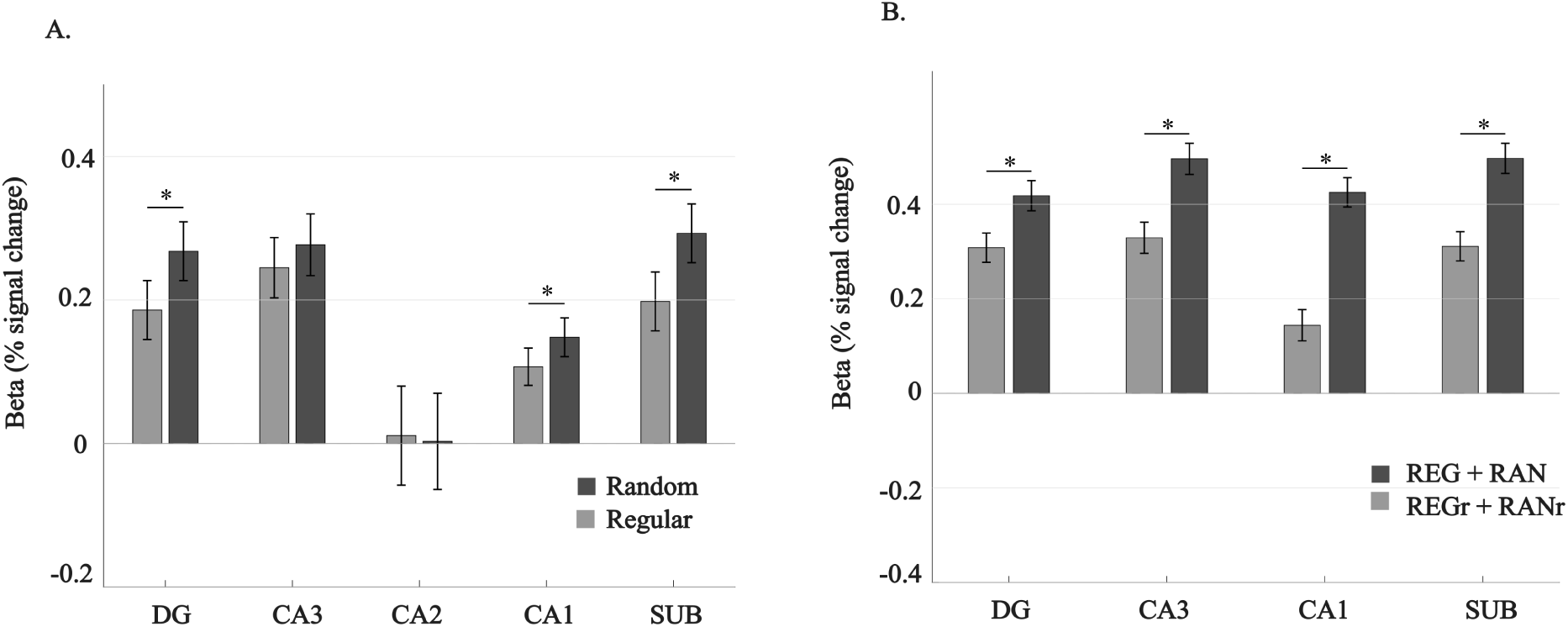
A. Main effects of regularity ([REGn+REGr>RAN+RANr]) in each subfield of the hippocampus. Light and dark grey bars represent the estimate of the responses to regular and random sounds, respectively (beta values obtained from the GLM analysis [percent signal change] - error bars represent the standard errors). The significant main effects are highlighted by a horizontal line on top of the bars and a star indicating q(FDR) <.05). B. Main effects of reoccurrence ([REGr+RANr>REGn+RAN]) in each subfield of the hippocampus. Light gray bars represent the estimate of the responses to reoccurring sounds (REGr and RANr) and dark gray bars the estimate of the response to novel sounds (REG and RAN). All other conventions are as in A.

For the memory effect, we first considered the interaction of reoccurrence and regularity by comparing the difference of reoccurring regular patterns and novel regular patterns (REGr vs. REGn) to the difference of reoccurring random patterns and novel random patterns (RANr vs. RAN) but found no significant effects in any of the subregions (for all statistical analyses see Supplementary figure S1). We therefore tested for the main effect of reoccurrence by contrasting reoccurring (REGr and RANr) to novel (REGn and RAN) patterns. This analysis revealed significant effects in all hippocampal subregions except CA2 (Figure 6B and supplementary S1), with lower activity for reoccurring compared to novel patterns.

## Discussion

Using UHF fMRI, we investigated the neural mechanisms supporting the implicit memory for reoccurring regularities (REGr) embedded in stochastic auditory sequences, which has been shown at the behavioural level [17]. Using the reoccurrence of five regular patterns, we replicated a RT advantage for reoccurring (REGr) versus novel (REGn) regularities (see Figure 2B), consistent with the gradual emergence of implicit memory [17, 22]. Immediately following the behavioral session, we collected fMRI responses from passively listening participants(T2* weighted GE-BOLD, 1.6 mm isotropic). The present study thus primarily captures post-learning neural responses. We replicated the larger responses for regular sounds compared to random patterns in a network of areas including bilateral inferior frontal gyrus (IFG) and superior temporal gyrus (STG - Figure 4 and [3, 6, 7]). Importantly, we extend these results in three ways. First, we observed striatal engagement in the processing of regular sounds (Figure 4), a region not previously associated with auditory regularity processing. Second, while the cortical regions (IFG/STG) and striatum showed increased responses to regularity compared to random sequences, hippocampal BOLD responses decreased (Figure 6). Third, a similar neocortical network was active when a reoccurring regularity was presented (Figure 5 and Figure 7D). Together, these findings highlight that the same core network of neocortical areas involved in the sensory processing of regularities [3, 6, 7] also supports their implicit memorization.

**Figure 7:**
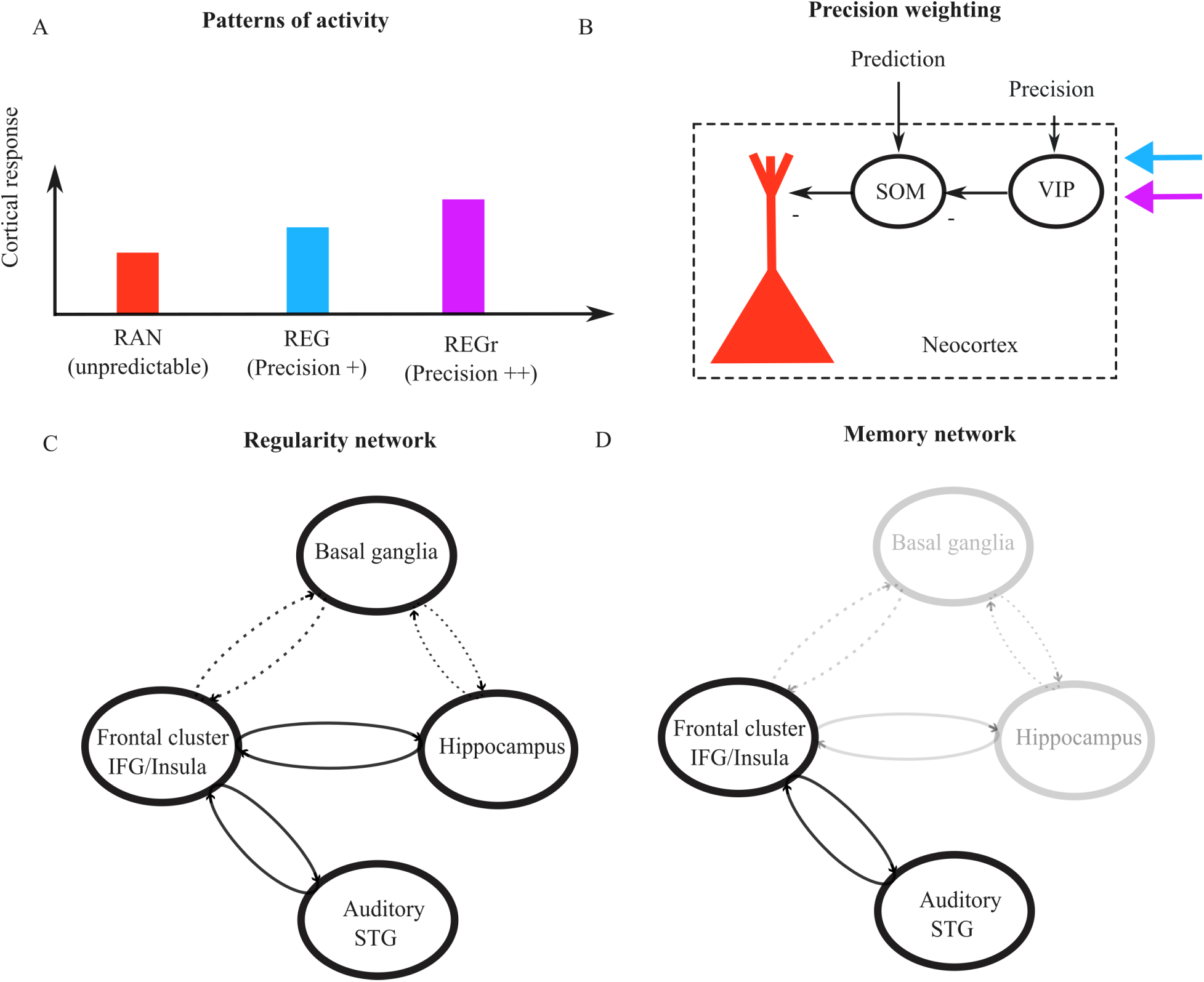
A. In our study, Regular patterns (blue) showed increased cortical activity over random patterns (red) and reoccurring regular patterns showed an even larger cortical effect(purple). B. In predictive coding frameworks, when stimuli are predictable, top-down predictions arising from higher cortical areas are believed to suppress activity in lower cortical regions. This effect is thought to arise from deep layers in higher cortical areas and targets superficial layers in lower cortical areas [10, 58]. The suppressive effect may occur via long-range GABAergic projections to the neocortex, or by activating local inhibitory interneurons (for example, somatostatin-positive (SOM+) interneurons) in superficial cortical layers targeting pyramidal neurons (here shown in red - for review see [10]). Precision on the other hand might counteract the suppressive effects of descending first-order predictions. Disinhibitory mechanisms often rely on vasoactive intestinal peptide-positive (VIP+) interneurons, which mediate disinhibition by inhibiting parvalbumin-positive (PV+) and/or SOM+ interneurons — cell types that typically suppress excitatory neuron activity [59]. The blue arrow represents increased precision reflecting the detection of any regularity, the purple arrow represents further enhanced precision due to the detection of a reoccurring regularity. Direct hippocampal projections might reach these inhibitory interneurons, or information might flow down from higher (frontal) to lower (temporal) level cortical areas. C. Depiction of the network involved in regularity detection. D. Network involved in the memory effect.

### A Bilateral IFG-STG network supports the detection of regularity

The increased activation in bilateral IFG and STG upon hearing a regularity replicates prior fMRI, MEG [3, 6, 7] and EEG [18] findings and is in line with broader evidence that this network underlies the processing of structure in unfolding sounds [25] and detection of violations [9, 26]. Our results align with studies demonstrating that detecting statistical regularities in complex auditory patterns enhances sensory responses (see Figure 7A and [3, 6, 7, 18, 27–31]). This stands in contrast with classical predictive-suppression accounts, which propose that predictable sensory inputs are “explained away” via top-down predictions, leading to attenuated cortical responses to expected stimuli [8, 9]. At first glance, one might attribute the increase in cortical activity to regular patterns to heightened arousal due to the salience of a regular pattern within a stream of random tones. However, this explanation is unlikely, as pupil size, an index of arousal, has been shown to decrease rather than increase upon the detection of regularities [32, 33]. A more likely explanation within the same predictive coding framework is that the increase in response to regular patterns reflects an increase in the precision of priors associated with these stimuli [26, 34, 35]. At the cortical level, precision modulation could be signaled by disinhibition of local cortical populations [10] (see Figure 7B). The local cortical circuit responsible for inhibiting ascending cortical activity, is in fact also capable of disinhibiting (i.e., the inhibition of inhibitory neurons) through vasoactive intestinal peptide-positive (VIP+ - Figure 7B), counteracting the suppressive effects of descending first-order predictions [36, 37]. We suggest that, for regularity alone, reduced uncertainty (i.e., increased precision of priors) may be increasing this disinhibitory input (from higher to lower regions), resulting in a larger response to regular compared to random sequences [38–40] (see Figure 7A).

### The putamen is implicated in the automatic discovery of structure in rapid sound sequences

In addition to the cortical network previously identified, we observed increased activation for regular sound patterns in the putamen, a subregion of the striatum within the basal ganglia (Figure 4). This finding extends earlier work by implicating subcortical learning systems in the processing of auditory regularities. The basal ganglia have long been considered central to sequence and habit learning, particularly in motor [41–43], habitual [44], and instrumental domains [45–48]. Their activity is modulated by learning in both rats [49, 50] and macaque monkeys [51], and accumulating evidence suggests a similar role in auditory learning processes, including auditory category learning [52], linguistic sequencing [53, 54], syntactic processing [55], and frequency discrimination in mice [56]. Computational accounts further propose that the basal ganglia support the gradual extraction of statistical regularities and the gating of cortical representations [57]. Thus, the putaminal involvement in auditory regularity processing we show here suggests that the basal ganglia contribute not only to motor and action-based learning, but also to the detection of structured sensory patterns.

### Hippocampus exhibits decreased activity for regular sequences

We also found significant hippocampal involvement in processing regular tonal patterns, though with decreased activity for regular compared to random sequences. Leveraging the higher specificity and spatial resolution afforded by UHF fMRI, we observed effects of regularity in the dentate gyrus, CA1, and subiculum—regions previously implicated in pattern separation ([60, 61]), a process through which the hippocampus transforms overlapping inputs into distinct, non-overlapping representations [62].

Interestingly, in the hippocampus all differential responses(i.e. both the main effect for regularity and reoccurrence) were found to be negative compared to the positive neocortical effects for regularity. Previous studies ([3, 6, 7]) demonstrated heightened ongoing MEG activity during the processing of regularity in the hippocampus. A potential reason for this difference in polarity of the effect (between MEG and fMRI) may be that linking electrophysiological activity to BOLD signals in the hippocampus remains fraught with inconsistency [63] — and this variability may be compounded by the region’s anatomical and functional complexity. Additionally, due to the temporal averaging inherent in the fMRI signal, it is possible that a transient initial increase in activity at the onset of the regularity (which has been the focus of previous MEG source analyses) is not captured by our data.

While the observed decrease in hippocampal activity upon hearing a regular pattern might initially seem counter-intuitive, this finding is consistent with how hippocampal function is thought to switch from errors to predictions during the learning process [64]. While early in the learning process the hippocampus up weights errors through dampening of predicted inputs, after learning responses to predicted inputs are thought to be sharpened, leading to increased discriminability but reduced response amplitudes at the population level. Thus, sharpening of neural population responses could explain the reduced BOLD activity we observe in the hippocampus for the regularity in general.

### Implicit auditory memory activates the same neo-cortical network as implicated in pattern discovery

By investigating the interaction of reoccurrence and regularity (i.e. whether the response difference between reoccurring and novel patterns was different for regular or random patterns) we aimed to identify genuine memory effects associated with the previously learned regularities and distinguish them from effects associated with increased signal to noise (SNR) due to the repetition of (any) stimulus. We observed a significant interaction (i.e. stronger reoccurrence effect for regular patterns than for random ones) within a similar neocortical network as the one found for the processing of regularity. Within the cortical microcircuit for predictive coding described earlier, the stronger response to REGr stimuli could be explained by increased precision of REGr patterns (compared to REGn RAN and RANr) during behavioral memory induction (Figure7B). This is consistent with the prediction-by-partial-matching model, which links stronger memory traces to increased precision estimates [17]. It should be noted that, on the group average surface, the memory effect was associated to a smaller frontal cluster (Figure7D) along the inferior bank of the inferior frontal gyrus and neighboring insular regions. While this may reflect the sampling of volumetric statistical maps (computed in a common [Talairach] space) to the cortical surface, the involvement of anterior insular regions, may be indicative of the strengthening of sensory representations [1]. Collecting more data from individual subjects and performing the analysis in native space would enable a more reliable assessment of insular cortex involvement.

### No hippocampal contribution to implicit auditory memory

In contrast, no hippocampal subregion showed an interaction between regularity and reoccurrence, indicating that the hippocampus did not differentiate between previously learned and novel reoccurrences. This pattern may indicate that the hippocampus contributes to the initial detection of structure and subsequently transfers this information to cortical networks where memory traces are then stored. This interpretation aligns with MEG findings showing hippocampal involvement during the early stages of regularity discovery, but not during the sustained phase of processing [3, 7]. We observed a general decrease in hippocampal activity for all reoccurring sequences, regardless of whether they were regular or random. Sharpening cannot easily explain this effect, unless the repetitions of RANr within the scanning session are sufficient to induce some form of rapid online learning of emerging reoccurrences. This pattern suggests that, rather than encoding the specific identity of reoccurring regularities, the hippocampus signals the degree of novelty in the auditory input. The fact that this decrease was observed across all subregions of the trisynaptic pathway (dentate gyrus, CA3, and CA1) may indicate the engagement of pattern completion processes, whereby reoccurring input patterns are efficiently reinstated from partial cues [11, 12, 65, 66]. Note that while the overall difference between reoccurring (REGr and RANr) and novel (RAN and REG) patterns may reflect effects associated with increased signal to noise (SNR) due to the repetition of (any) stimulus - an account that might also explain earlier findings of hippocampal involvement in reoccurring stimuli [14], this interpretation, in our case, is challenged by the observation of reduced responses to the reoccurring stimuli. We also tested for a main effect of reoccurrence in the cortex but found no significant activation. Together, this pattern of responses (interaction of reoccurrence and regularity in neocortex and main effect of reoccurrence in hippocampus) may imply that while the hippocampus broadly signals repetition, cortical regions may selectively use this information to enhance predictions or representations of structure only when the environment is regular and thus predictable.

Given that participants were extensively exposed to the regularities prior to scanning, our study primarily captures neural responses after learning has taken place. Within this post-learning context, both novel and reoccurring regularities activated a well characterized auditory and frontal network, replicating prior findings [3]. Interestingly, while cortical regions such as frontal and auditory regions showed an amplification of the difference between reoccurring regular patterns and regular patters when compared to the difference of reoccurring random and random patterns, the hippocampus exhibited a main effect of reoccurrence and regularity with reduced activity to both regular (compared to random) and reoccurring (compared to novel) patterns.

The finding that the memory effect is localized to the cortex, specifically within a frontal cluster and superior temporal gyrus (STG), is consistent with the notion that memory may be stored through potentiated synaptic weights within the same sensory regions that initially process auditory input, rather than in a distinct memory store. Although null results must be interpreted cautiously, the absence of hippocampal involvement in the memory effect aligns with the idea that the hippocampus contributes primarily to the short-term discovery of structure, subsequently transferring this information to cortical regions for longer-term storage. This interpretation is further supported by findings in aging [22], which show that while older adults exhibit a reduced memory advantage (indexed by reaction times) compared to younger adults, potentially reflecting hippocampal differences, the persistence of memory over time remains intact, with no evidence of decay. This pattern may also help explain why auditory memory tends to be relatively preserved in dementia.

Future work could investigate the dynamic process of learning, focusing on how the identified network evolves over time or changes after sleep, rather than examining only the end state of consolidated learning. Additionally, examining the layer-specific responses within this network may provide deeper insight into the mechanisms underlying different stages of learning.

## Methods

### Participants

Fourteen participants (13 women), aged 21 to 30 participated in the study. All participants provided informed consent prior to participation and received either monetary compensation or academic credit for their involvement. Exclusion criteria included any known neurological or psychological disorders, impaired hearing abilities, and any conditions that contraindicated MRI procedures, such as the presence of metal objects in the body. The study was approved by the ethical review committee of the Faculty of Psychology and Neuroscience of Maastricht University, following the principles expressed in the Declaration of Helsinki. The MRI data were collected at the Scannexus facility of Maastricht University (Maastricht, The Netherlands).

### Behaviour

#### Stimuli

We used stimuli similar to those in [17]. Stimuli were five second sequences composed by 50 ms tone pips of different frequencies (20) sampled in the range between 222 Hz and 2000 Hz. The order in which the frequencies were successively presented defined the different conditions that were otherwise identical in their spectral and timing profiles. Sequences were either random throughout (RAN; tones uniformly sampled from a pool of 20 values) or transitioned into a regular structure (REG), which consisted of a repeating 20 tone (1 s long cycle) pattern. The transition occurred between 1500 and 2000 ms after sequence onset such that each sequence contained between 3 and 3.5 REG cycles. Unbeknownst to the participants, a 5 REG patterns - unique to each individual - reoccurred (repeated regularly) such that REG sequences were either novel (REGn) or reoccurring patterns (REGr). The stimulus set also included control stimuli (STEP and CONT) with which we estimated listeners’ basic reaction time to a simple sound change. To estimate the regularity detection latency, we subtracted response times (RTs) to STEP from RTs to the REG conditions. Each participant completed five blocks, each lasting approximately eight minutes.

#### Behavioural session

Each trial consisted of the presentation of one sequence (from one of the 4 possible conditions - see Figure 1). Participants were familiarized with the stimuli and the task (detecting frequency change or regularity) by listening to twelve trials (5 RANREG, 5 RAN, 2 STEP, 1 CONT) prior to the first behavioural block. Within each block, participants listened to 100 trials with an inter trial interval of 1000ms. Participants were asked to monitor transitions by pressing a space bar as soon as they heard a regularity occurring, or when a change in frequency occurred. To ensure engagement and optimize reaction times (RT), feedback on response accuracy and speed was provided at the end of each sequence. Given the importance of RT as a key performance metric in these experiments, participants were explicitly encouraged to respond as quickly as possible. The feedback procedure followed a previously established approach [3]). A green circle was displayed if the response occurred within 2200 ms of the onset of regularity in RANREG sequences or within 300 ms of the frequency change in the STEP condition. For slower responses, an orange circle was shown for RTs between 2200 and 2600 ms in RANREG trials and between 300 and 600 ms in STEP trials. Responses exceeding these limits were indicated by a red circle. Participants were instructed to aim for as much ‘green’ or ‘orange’ feedback as possible. Each block lasted approximately eight minutes containing 30 REGr sequences (five unique sequences reoccurring six times), 30 REGn, 30 RAN, 5 STEP, and 5 CON sequences. Each participant completed five blocks, and blocks were separated by brief breaks. Auditory stimuli were presented using PsychTool-Box in MATLAB (version 9.2.0, R2017a) in an acoustically shielded environment. Sound levels were set to a comfortable intensity and adjusted individually by each participant.

#### Analysis

To assess participants’ sensitivity to the task, we calculated d prime for each block, providing a measure of their ability to discriminate between regular and random sequences while controlling for response bias. Response times were analysed using a repeated measures ANOVA with within-subject factors Block and Condition (REGn and REGr). To approximate the lower bound of the processing time required to detect a frequency transition, we subtracted reaction times (RTs) to the STEP condition from RTs to the REGn and REGr conditions. This adjustment provides a corrected RT measure, isolating the time attributed to detecting the regularity from basic auditory processing and motor response latency.

### fMRI session

#### MRI data acquisition

Anatomical (T1 weighted) images were acquired on a 7-T Siemens MAGNETOM MRI scanner using a Magnetization Prepared 2 Rapid Acquisition Gradient Echo (MP2RAGE) sequence [67] (240 contiguous slices per volume; Repetition Time (TR) = 1000 ms; Echo Time (TE) = 2.47 ms; matrix size: 320 × 320; slice thickness: 0.7 mm; flip angle = 5 degrees; 0.7 mm isotropic resolution with a field of view (FoV) of 223 × 223 mm). Functional images (T2* weighted) were acquired with multiband accelerated echoplanar imaging (EPI) sequences [68] (TR = 1000 ms; TE = 20 ms; GRAPPA = 2; MultiBand factor = 3; FoV = 186 x 186 mm; resolution = 1.6 mm isotropic). Each participant completed six runs, resulting in the collection of a total of 2,730 volumes, each run lasting approximately 10 minutes. Five additional volumes with reversed phase encode polarity were also acquired for correction of geometric distortions after the collection of the anatomical data and before the first experimental run.

#### Stimuli

The same stimuli used in the behavioural session were passively presented during scanning as a continuous stream of random tones (RAN) with occasional embedded bursts of regular (REG), reoccurring regular (REGr), or reoccurring random (RANr) patterns. RANr patterns were added to control for the mere repetition effect, and consisted of reoccurring 3 second long RAN sequences that were presented the same number of times as REGr (different for each participant). STEP and CONT sequences were no longer presented in the scanner. Bursts of REGn, RANr, and REGr were always 2000ms long - ensuring two full cycles of e.g. the REG component. Each run started with ten seconds of rest and consisted of two parts (Part 1 and Part 2) separated by twenty seconds of rest. After Part 2, we always acquired an additional twenty seconds of rest. Each part consisted of forty sequences concatenated with no silences (5 REGr (identical to those in the behavioural session), 5 REGn (novel in each run), 5 RANr sequences, and 25 RAN sequences. Our original intention was for Part 2 to include always novel REG and RANr subsequences. However, due to an error in the stimulus presentation code, the subsequences used in Part 2 of each run (5 REGr, 5 REG, and 5 RANr) were the same as those presented in Part 1, with different RAN sections preceding them. As a result, each REGr stimulus was repeated twelve times over the scanning session (two times per run), and the same was true for the RANr stimuli. REGn stimuli were presented two times per run each (but not repeated across runs), while RAN stimuli were never repeated.

#### Anatomical and functional data analysis

BrainVoyager (v22.2, Brain Innovation, Maastricht, The Netherlands) was used in all anatomical and functional pre-processing steps unless otherwise specified. The unbiased T1 weighted image obtained from the MP2RAGE was de-noised [69] and the skull and dura were removed using Nighres [70]. We then corrected for inhomogeneities and up-sampled the anatomical volumes to 0.8 mm isotropic. The anatomical data data were then projected to Talairach space and the white matter - gray matter boundary was obtained using the advanced segmentation pipeline in BrainVoyager. The surfaces representing the left and right hemispheres of all individuals were used for cortex based alignment [71], and the average cortical surface across individuals was used for visualization purposes. The anatomical data in native space were used to define hippocampus regions of interest using the Hippocampus Segmentation Factory (HSF)[24]. We divided the hippocampus into five subfields, Subiculum, Dentate Gyrus, and Cornu Ammonis areas 1-3 (CA 1-3).

Functional data were slice scan time corrected (sinc interpolation), motion corrected (sinc interpolation), and high pass filtered (removing frequencies lower than two cycles per run). After visual inspection, the data of three participants required additional correction for distortions between runs (i.e. after motion correction there were noticeable distortions across runs due to excessive motion between runs). To correct for these distortions, we applied non-linear alignment using Advanced Normalization Tools (ANTs). All data (all subjects and runs) were then corrected for distortions caused by phase encoding using the reversed phase polarity acquisitions and Topup (FSL, v6.0.4). Finally, functional data were aligned to the upsampled anatomical images in native space using boundary-based registration. We ten resampled the functional data in the anatomical space at a resolution of 1.6 mm isotropic (sinc interpolation) and obtained two sets of functional data per volunteer, one in the original anatomical space and one in Talairach space. All statistical analyses were performed using a General Linear Model (GlM - fixed effects) in which the transitions (to REG, REGr, RAN and RANr) were coded as separate predictors together with the presentation of tone pips representing the first three seconds of every sequence (RANon). We separated the predictors for Part 1 and Part 2 of every run (to consider the potential effects of immediate repetitions within a run). All predictors were convolved with a standard (double gamma) haemodynamic response function. For the analysis of the cortical responses, the GLM was fitted to time series normalized in Talairach space and concatenated across subjects. The statistical analyses of the hippocampus were computed using a ROI approach, in which the time course of each ROI (in the original space) was first obtained by averaging all voxels (separately for each ROI) and then concatenated across subjects (for the fixed effect group analysis). All statistical tests were corrected for multiple comparisons using False Discovery Rate (FDR). We tested for regularity by contrasting the responses to regular (REG+REGr) and random (RAN+RANr) subsequences, and for regularity by reoccurrence interaction by comparing (REGr - REG) with (RANr - RAN). Prior to running both analyses on the data collapsed across Part 1 and Part 2, we verified that there were no significant interactions with part and in absence of interactions we report the main effects (i.e. the effects averaged across part 1 and 2).

## Supplementary Materials

### Individual responses to sound

**Figure S1:**
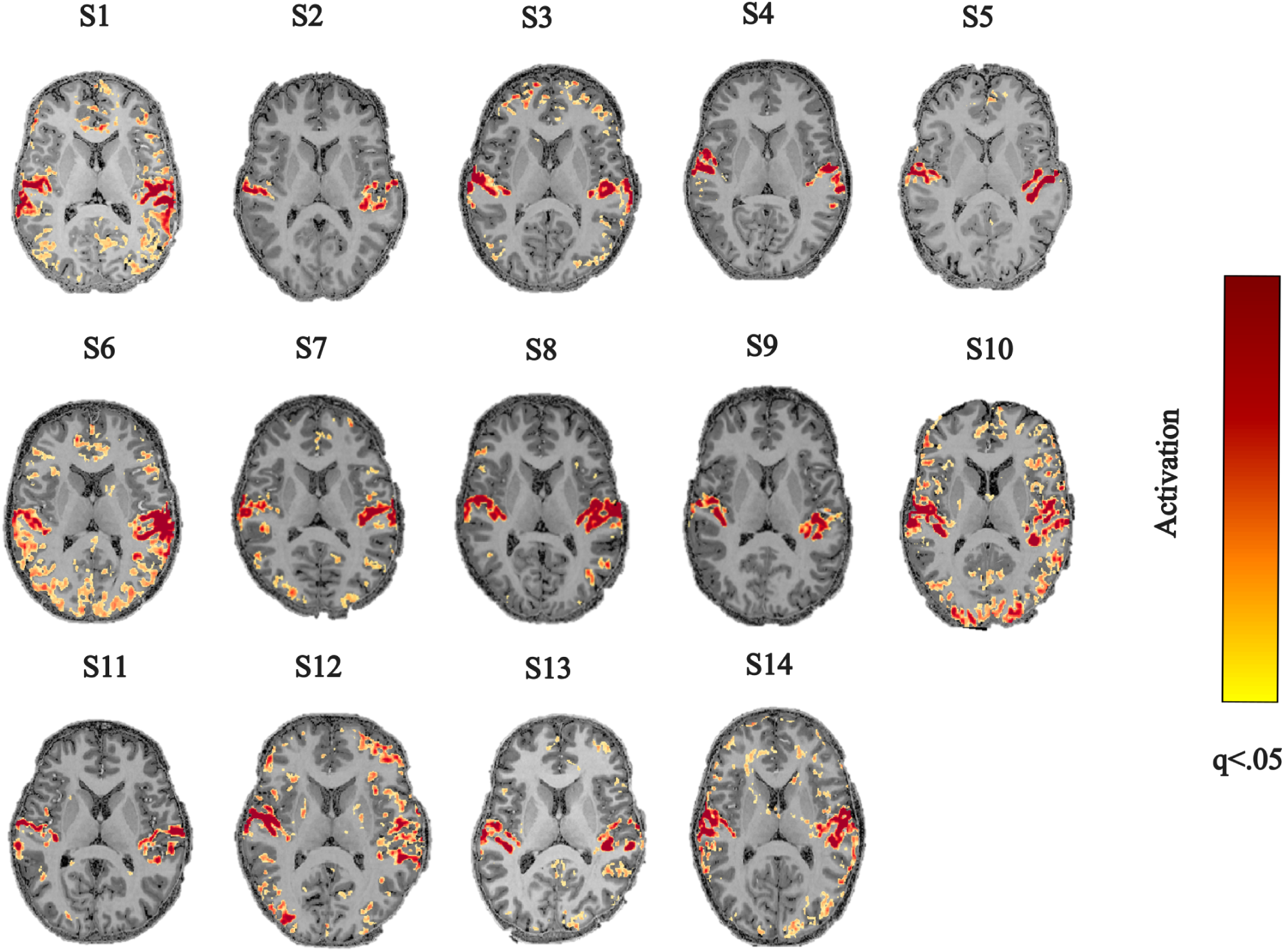
Individual statistical analysis of the response to any sound (RANREG, RAN-REGr, RANRAN, RANRANr) in our design compared to no sound presentation (t-omnibus test). The results are presented on a transversal slice cutting through bilateral temporal areas. The statistical map was corrected for multiple comparisons using FDR and thresholded at a level of q(FDR)<0.05.

### Hippocampus segmentation

**Figure S2:**
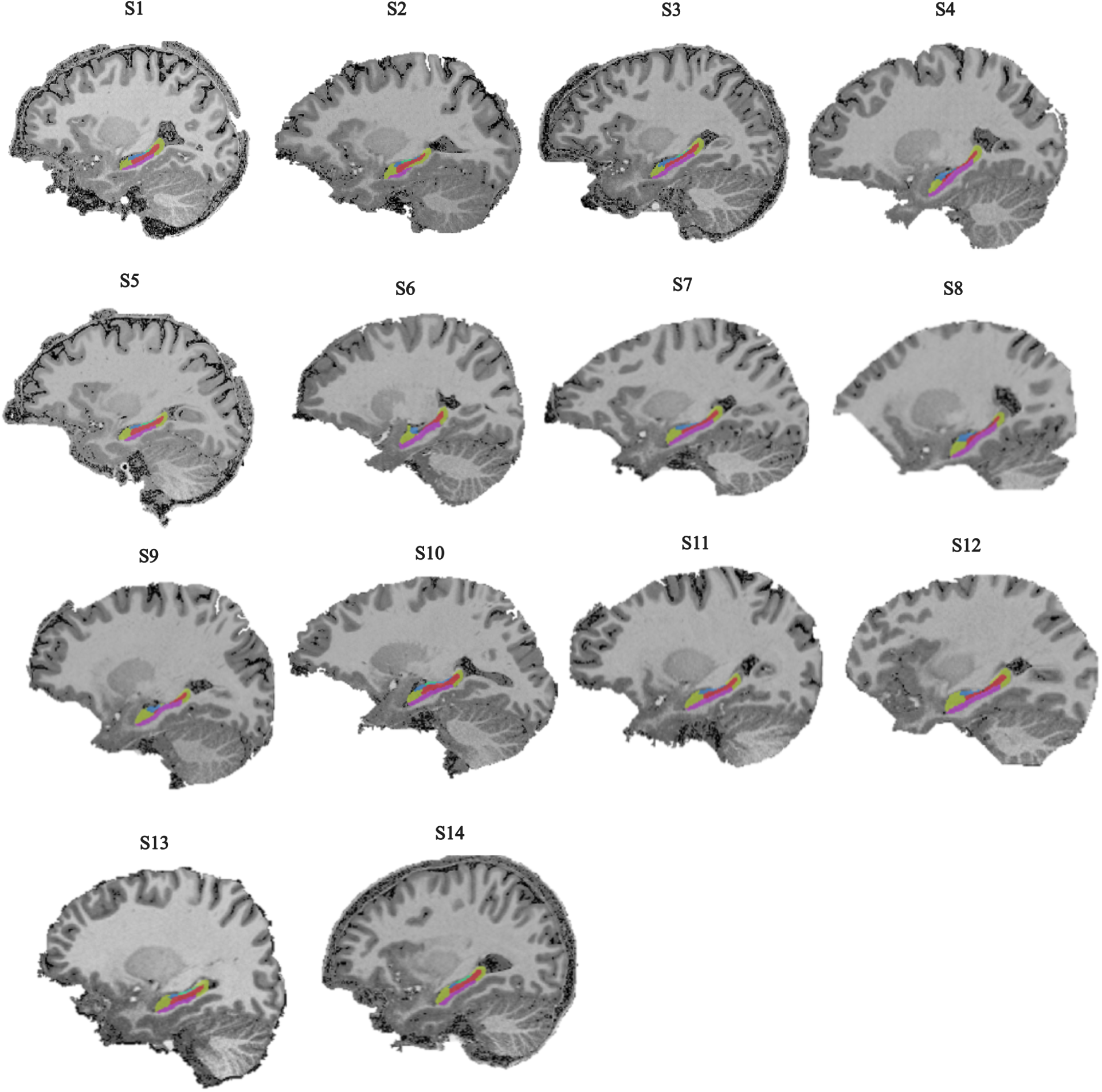
Individual segmentation of hippocampal subfields (one sagittal slice through the left hemisphere). The individual ROIS are overlaid in different colour transparency over the individual anatomical data, dentate gyrus, subiculum, cornu ammonis 1-3). Red color represents dentate gyrus, yellow is CA1, green is CA2, and purple is Subiculum

### Summary of test statistics across hippocampal subfields

**Table S1:**
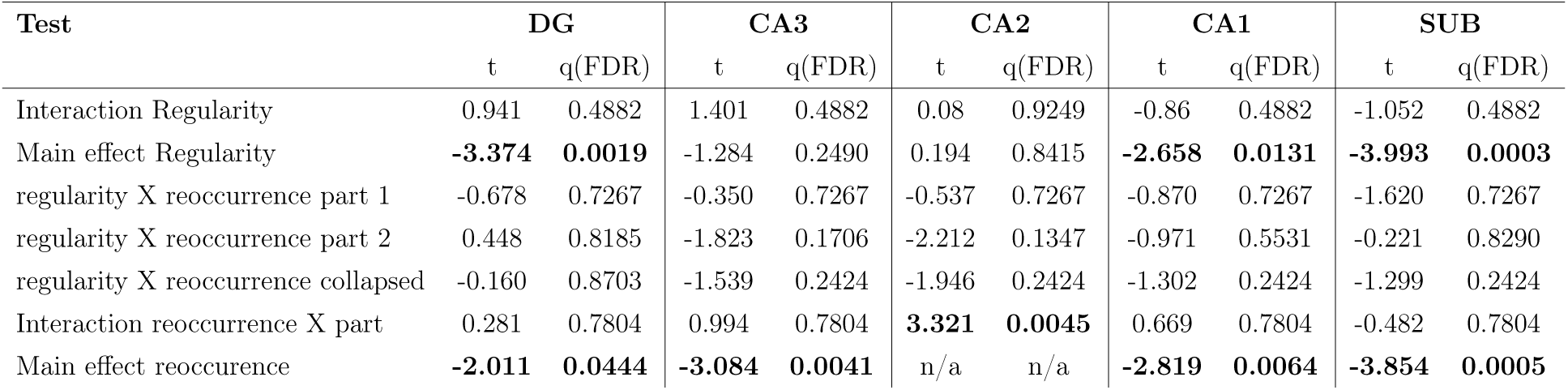
In the hippocampus we tested for the same interactions as in the neocortex. For the regularity effect ([RANREGr and RANREGn] versus [RANRANr and RANRAN]) we found no interaction in any of the subregions and therefore analyzed Part 1 and Part 2 together (main effect Regularity). For the regularity by reoccurrence interaction ([RAN-REGr - RANREG] versus [RANRANr - RANRAN]), we checked whether the interaction was present in Part 1 and Part 2 separately. After establishing this interaction was not present in the seperate parts, we tested whether the interaction was present when collapsed across the parts. In none of the subregions this was the case, hence we followed up by testing the main effect of reoccurrence ([RANREGr+RANRANr>RANREG+RANRAN]. We first tested whether this main effect differed between Parts. This was only the case in CA2. For all other subregions, we did not find the reoccurrence by part interaction and hence collapsed across the two Parts (main effect reoccurrence)

## Notes

### Competing Interest Statement

The authors have declared no competing interest.

